# Microplastics may reduce the efficiency of the biological carbon pump by decreasing the settling velocity and carbon content of marine snow

**DOI:** 10.1101/2023.06.23.545915

**Authors:** Cordelia Roberts, Clara M. Flintrop, Alexander Khachikyan, Jana Milucka, Colin B. Munn, Morten H. Iversen

## Abstract

Plastics are pervasive in marine ecosystems and ubiquitous in both shallow and deep oceans. Microfibers, amongst other microscopic plastics, accumulate in deep sea sediments at concentrations up to four orders of magnitude higher than in surface waters. This is at odds with the fact that most microfibers are positively buoyant; therefore, it is hypothesized that settling aggregates are vectors for downward transport of microfibers in the ocean. However, little is known about the impact of microfibers on carbon export. We formed diatom aggregates with differing concentrations of microfibers using roller tanks and observed that microfiber addition stimulated aggregate formation, but decreased their structural cohesion and caused them to break apart more readily, resulting in smaller average sizes. Incorporation of positively buoyant microfibers into settling aggregates reduced their size-specific sinking velocities proportional to the microfiber concentration. Slower sinking may extend aggregate retention time in the upper ocean, thereby increasing the time available for organic matter remineralization in the upper water column. Here, we show that microfiber concentrations typical of those in the English Channel and Atlantic Ocean decrease potential export flux by 15-50%. Present day microfiber concentrations in surface waters may therefore be substantially reducing the efficiency of the biological carbon pump relative to the pre-plastic era.

## Introduction

High production and usage of plastics in combination with poor waste management has resulted in enormous amounts of plastic litter in the oceans (Ryan and Moloney 1993; Thompson et al. 2004; Barnes et al. 2009; Jambeck et al. 2015). Increasing amounts of plastic debris in the marine environment are affecting marine biota and ecosystem functioning (Gregory 1996; Andrady and Neal 2009). Plastic particles with sizes between 0.1 μm and 5 mm, referred to as microplastics (Barnes et al. 2009), are of particular concern because of their pervasiveness in marine food webs (Setälä et al. 2018; Carbery et al. 2018). Microplastics originate from materials used in manufacturing, cosmetics, and washing machine effluent in the form of microfibers (“primary” microplastics) as well as from fragmentation of macroplastic debris (“secondary” microplastics) (Thompson et al. 2004; Ryan et al. 2009). Of these, synthetic microfibers from washing machine effluent constitute the single largest source of marine primary microplastics (Browne et al. 2011; Pirc et al. 2016).

Microfibers have been observed to accumulate in deep sea sediments at concentrations up to four orders of magnitude higher than in surface waters (Woodall et al. 2014) and are ingested by deep sea organisms (Taylor et al. 2016). This is at odds with the fact that most microfibers are positively buoyant (Kaiser et al. 2017; Porter et al. 2018) and it has therefore been hypothesized that microfibers reach the deep sea after incorporation into settling material such as marine snow (Van Cauwenberghe et al. 2013; Woodall et al. 2014). Marine snow, i.e. aggregated organic matter with diameters larger than 0.5 mm, form through the aggregation of smaller particles, typically diatoms, phyto-, and zoodetritus (Alldredge and Silver 1988; Alldredge and Gotschalk 1988). It is the settling of marine snow that drives the biological carbon pump (BCP) by exporting carbon fixed by phytoplankton from the surface to the deep ocean (Volk and Hoffert 1985). The efficiency of the biological carbon pump, i.e. the fraction of primary production that is exported to depth, is determined by the turnover and the settling velocity of sinking aggregates, e.g. marine snow, phytoplankton aggregates and cells, and zooplankton faecal pellets (Ploug et al. 2008; Iversen and Ploug 2010).

Microplastics have been shown to be efficiently incorporated into aggregates, leading to the downward transport of microplastics in the water column (Porter et al. 2018; Michels et al. 2018; Kvale et al. 2020; Galgani et al. 2022). At the same time, incorporation of microplastics into marine aggregates makes them more buoyant and causes the aggregates to sink slower than they would without microplastic, although to date, this has only been shown for microbeads (Long et al. 2015; Porter et al. 2018). Additional evidence suggests the same holds true for gelatinous zooplankton faecal pellets (e.g., salp pellets [Wieczorek et al. 2019]) and copepod fecal pellets (Shore et al. 2021). Microplastics have also been found in marine aggregates in situ (Zhao et al. 2018). Most studies to date have focused on the transport of microfibers, but not on the impact of microplastics on particulate organic carbon export. To improve predictions of the efficiency of the biological pump in a world with ever-increasing amounts of plastic (Geyer et al. 2017), studies are needed to assess the impact of microfibers on the settling velocity and carbon content of marine snow and the implications for carbon export. In this study, the ubiquitous diatom *Skeletonema marinoi* was incubated with three different concentrations of synthetic microfibers common in clothing fabrics to assess how microfibers influence aggregation dynamics, settling velocity, and export of carbon to the deep sea.

## Material and Methods

### Microfiber composition and phytoplankton cultures

We formed marine snow during roller tank incubations with diatoms (at constant concentration) and different concentrations of microfibers that were shorter than 3 mm. We determined the microfiber composition using a confocal Raman spectrometer (NTEGRA Spectra, Eindhoven, The Netherlands). The Raman spectra showed that the microfibers were primarily composed of dyed cotton and polyamide (Fig. S1). This was identified from comparisons to known Raman spectra of dyed cotton and polyamide (Lepot et al. 2008). Still, some peaks within the spectra were unmatched, suggesting that the microfiber contained additional materials.

Cultures of *Skeletonema marinoi* (Sarno & Zingone 2005) were grown in GF/F-filtered natural seawater with salinity 32 enriched with f/2 medium (Guillard 1975) and silicate at a 1:1 molar ratio of silicate to nitrate. Cultures were grown at 15°C under a 14:10 h light:dark cycle at 150 μmol photons m^−2^ s^−1^ until they reached the stationary growth phase at 5.85 × 10^7^ cells mL^-1^.

### Aggregate formation and settling velocity

Aggregates were formed by incubating *S. marinoi* cultures at a concentration of 5.4 × 10^4^ cells mL^-1^ in four 1.15 L roller tanks and rotating them at 3 rpm at 15°C under low light conditions (∼30 μmol photons m^−2^ s^−1^). The microfiber concentrations in the four roller tank treatments were representative of current oceanic concentrations:

i. low concentrations (“low”) of 240 (±0.2) microfibers L^-1^. This amount is close to concentrations found in the western English Channel (approx. 270 microplastics L^-1^, (Cole et al. 2014);
ii. medium concentrations (“medium”) of 680 (±0.9) microfibers L^-1^;
iii. high concentrations (“high”) of 840 (±1.1) microfibers L^-1^. This is in the range of concentrations found in the Atlantic Ocean (1150 microplastics L^-1^; Kanhai et al. 2017)
iv. no fibers (“control”) were added to the fourth roller tank, which acted as a control without microfibers.

At five time points over a period of 163 hours (24h, 48h, 118h, 143h and 163h), recordings of the rotating tanks were made using a commercial digital single-lens reflex camera equipped with standard lens of 50 mm focal length. This enabled detection of aggregates >0.5 mm for measuring aggregate formation, size distribution, abundance, and size-specific settling velocities for each treatment. Aggregate size was measured using the projected area of each aggregate and calculating the equivalent circular diameter (ECD). The video recordings were used to determine size-specific sinking velocities of the formed aggregates based on the method developed by Ploug et al. (2010). This method is based on the observation that aggregates follow circular trajectories around a center (x_n_) from which the settling velocity can be predicted knowing the distance between the aggregate orbit center to the center of the roller tank (x_b_). A circle radius (R_n_) was calculated for the center position of each aggregate using the equation:

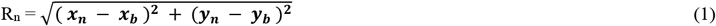

where ***x***_***n***_ and ***y***_***n***_ are the particle positions, n= 1, 2, …., N, and (***x***_***b***_, ***y***_***b***_) is the putative center of the aggregate orbit. Using the radii and the orbit center position, an idealised circle was plotted. The circle was manipulated to find the best fit of the actual aggregate orbit, R_n_, to the center position for each aggregate. The value produced for x_b_ was used to calculate the aggregate settling velocity using the equation:

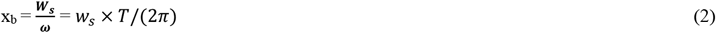

where w_s_ is the settling velocity of the aggregate (cm s^-1^), ω = 2π/T is the rotation rate (s^-1^) and T is the rotation period of the roller tank.

### Potential export flux

At the end of the experiment, aggregates of known volume from each of the four treatments were filtered onto 25 μm pre-weighed GF/F filters. Filters were dried at 40°C for 48h and re-weighed on an ultra-microbalance (UMX2, Mettler Toledo, USA) with a mass readability of ±0.1 μg to determine their volume-specific dry weight (DW). After fuming with hydrochloric acid, filters were measured with an elemental anlayzer isotope ratio mass spectrometry (EA-IRMS, ANCA-SL 20-20, Sercon Ltd. Crewe, UK) with a precision of ±0.7 μgC to determine the PON and POC content of aggregates. The potential export flux was estimated for each treatment as an average of the final three time points (118h, 143h and 163h) of the experimental period. This was done for each treatment by using the POC-to-volume ratio to determine the POC content per aggregate and multiplying with the measured settling velocity to get potential POC flux from each aggregate:

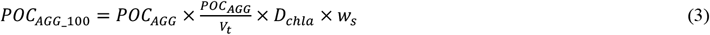

where POC_AGG_ is the POC content of each aggregate (μgC agg^-1^), V_t_ is the tank volume (ml), D_chla_ is the export distance from the fluorescence maximum to 100 m depth (here assumed to be 80 m) and w_s_ is the aggregate sinking velocity (m d^-1^). The potential POC flux was summed for all aggregates in each treatment to get total potential export flux using the equation:

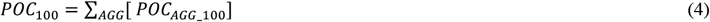

where POC_100_ is the estimated POC flux to 100 m for each microfiber concentration (gC m^-2^ d^-1^).

## Results

### Aggregate formation and structure

We incubated the diatom *Skeletonema marinoi* in roller tanks without and with added microfibers at three different concentrations: low (240 ±0.2 L^-1^), medium (680±0.9 L^-1^), and high (840±1.1 L^-1^). The low and high concentrations were representative of microfiber concentrations observed in the English Channel and the Atlantic Ocean, respectively. There was a positive linear correlation between initial microfiber concentrations in each treatment and the number of microfibers counted in aggregates at the end of the incubation (Fig. 3a). During the first 48 hours of roller tank incubation, aggregate formation followed a similar pattern in all treatments, with 7-10 comparably sized aggregates formed (Fig. 1). Between 48 – 118 hours, aggregation dynamics started to diverge among treatments. At zero, low, and medium microfiber concentrations, both the number and size of aggregates increased. At high microfiber concentration, average aggregate size also increased but the number of aggregates remained low. After 118 hours, there was a continued increase in larger aggregates accompanied by a decrease of smaller size classes in the treatment with no added microfibers. This effect was not as pronounced in the treatments with added microfibers, especially at medium and high concentrations.

**Fig. 1.**
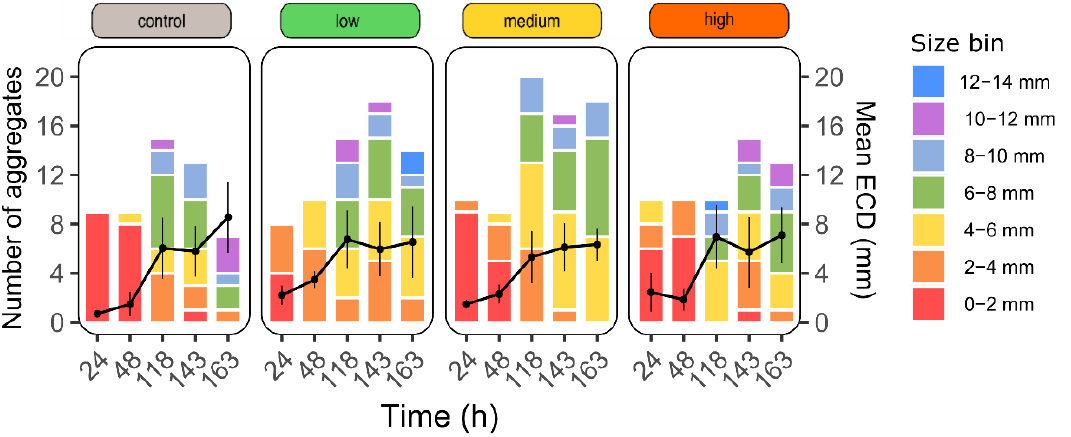
Number of aggregates in each size bin (primary y-axis) and mean equivalent circular diameter of aggregates (black line ± standard deviation) (secondary y-axis) throughout the study for each treatment: without addition of microfibers (control) and for low, medium, and high amounts of microfibers added to the diatom (*S. marinoi*) incubations.

### Aggregate settling velocity

We determined the size-specific sinking velocities of all aggregates in the different treatments at five time points during the incubations: after 24h, 48h, 118h, 143h, and 163h. Over the course of the experiment, some aggregates were observed to be positively buoyant, i.e., they were rising rather than sinking as indicated by their negative calculated sinking velocities. Buoyant aggregates were only observed in treatments to which microfibers were added, and overall, the number of positively buoyant aggregates decreased over time (Fig. 2): after 24 hours of roller tank incubation, 22% of the aggregates in the microfiber treatments were rising with a mean velocity of 6.0 (±2.8) m d^-1^. This decreased to only 10% of the aggregates at 48 hours (10.9 ± 5.2 m d^-1^) and at 118 hours none of the aggregates were rising. After 143 hours, 10% of the aggregates in the microfiber treatments were rising with a mean velocity of 3.8 (±1.85) m d^-1^ (Fig. 2).

A clear positive relationship between aggregate size and settling velocity could be observed after 48h (Fig. 2). Due to aggregate fractality, their size-to-settling relationship follows a power law function, but in the range observed in this study the settling velocity increased almost linearly with size. Generally, size-specific settling velocity was lower for the microfiber treatments compared to the control treatment without added microfibers (Fig. 2). After 143h, the size-specific sinking velocities decreased with increasing microfiber concentrations (Fig. 2).

**Fig. 2.**
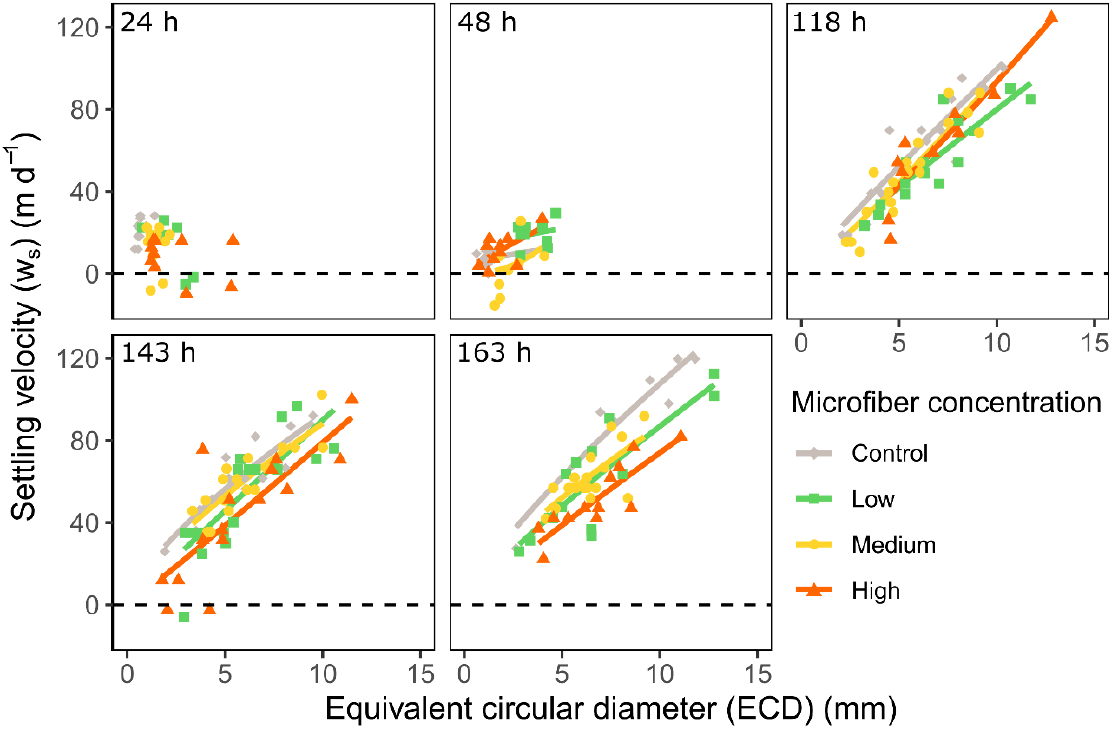
Aggregate settling velocities (w_s_) plotted against their equivalent circular diameter (ECD) throughout the study.

### Potential export flux

Total aggregate volume did not significantly differ between treatments (ANOVA p = 0.1207) (Fig. 3b). However, particulate organic carbon (POC) and particulate organic nitrogen (PON) content of individual aggregates decreased with increasing microfiber concentrations (Fig. 3c, Table S1). Combined, the decrease in carbon content and decrease in size-specific settling velocity in the microfiber treatments resulted in a 15-50% reduction in potential carbon flux compared to the control (Fig. 3d, Table S1), although the difference was not statistically significant due to high variability in potential C flux, especially in the control treatment (Table S2).

**Fig. 3.**
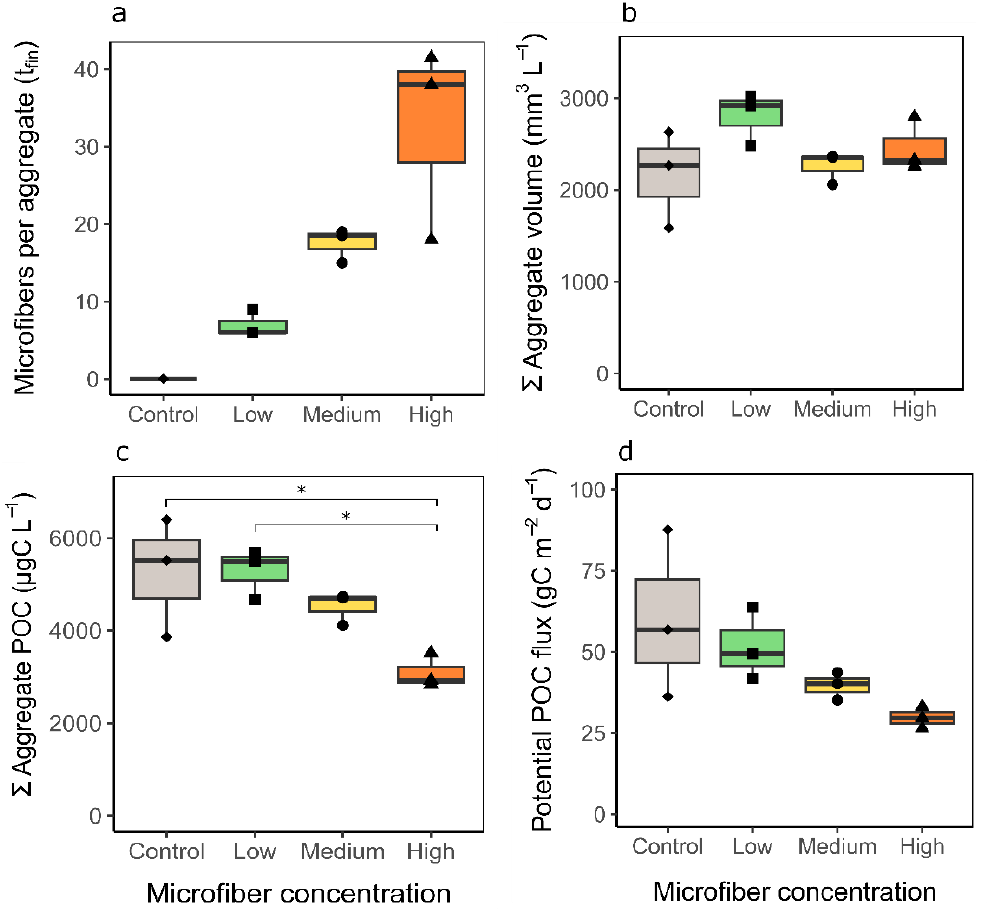
Microfiber content per aggregate (a), total aggregate volume (b), total aggregated POC (c), and potential POC flux to 100 m (d) as a function of microfiber concentration at t_0_ for aggregates averaged from days 5-7 for each treatment. An asterisk (*) denotes a statistically significant difference (Table S2).

However, there was a statistically significant difference in the number of microfibers incorporated into aggregates and the magnitude of potential C flux (Fig. S2). The relationship between the number of incorporated microfibers and potential C flux (g C m^-2^ d^-1^) was best described using an exponential regression:

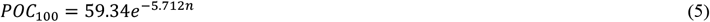

where n is the number of microfibers incorporated per aggregate volume (mm^-3^), R^2^=0.97.

## Discussion

The production of marine snow in surface waters and its subsequent transport to depth via the biological carbon pump plays a crucial role in the drawdown of natural and—increasingly abundant—anthropogenic CO_2_ (Wilson et al. 2022). It has previously been suggested that marine aggregates are vehicles for vertical transport of buoyant microplastics to the seafloor, (Woodall et al. 2014; Long et al. 2015; Porter et al. 2018). To date, most studies have focused on microbeads as a model plastic pollutant, despite microfibers being the most common type of microplastic identified in the ocean (Liu et al. 2022, and references therein). Here, we not only showed that marine aggregates incorporate plastic microfibers and can transport them to depth, we also measured the direct effects of microfiber incorporation on the formation, sinking velocity, and carbon content of marine snow, and discuss potential impacts on the efficiency of the biological carbon pump and wider marine carbon cycling.

Additions of microfibers to diatom cultures in roller tanks resulted in formation of aggregates that contained substantial numbers of microfibers, proportional to the initial microfiber concentration in the treatment. Incorporation of microfibers into aggregates had a measurable impact on key aggregate properties, such as size and settling velocities. Diatom cultures with added microfibers produced more, smaller, slower sinking aggregates than aggregates in the control treatment, impacts of which have the potential to decrease the magnitude of export flux.

In general, smaller aggregates sink slower than larger aggregates of a similar composition, but this was not the only reason for the observed lower sinking velocities within microfiber treatments: size-specific sinking velocities also decreased as a function of increasing microfiber concentration. Microfibers can decrease the setting velocity of artificial aggregates, in comparison to artificial aggregates without microfibers and with alternative microplastics (Porter et al. 2018). Our results indicate that the decrease in settling velocities observed in the microfiber treatments was due to increased buoyancy of the aggregates. In fact, during the first phase of the study we observed positively buoyant, non-sinking aggregates, suggesting microfibers within aggregates may delay export of organic material until aggregate mass density increases and aggregates start to sink. Evidence of buoyant particles and reduced sinking velocity seemingly contradicts hypotheses of marine snow as a vector of microplastics to depth. As export of marine snow is not limited to gravitational sinking, additional mechanisms transporting microplastics to depth may include particle injection pumps associated with the biological carbon pump, such as the eddy subduction pump, mixed layer pump and the mesopelagic migrant pump (via zooplankton grazing and defecation at depth (Boyd et al. 2019). Additionally, microbial communities associated with microfibers (Zettler et al. 2013; Vaksmaa et al. 2022) may also influence the sinking of microplastics (Amaral-Zettler et al. 2021) and can induce rapid aggregation of biogenic particles (Michels et al. 2018) which likely extends to microplastics. To determine how microbial colonization of microplastics influences aggregate dynamics, further laboratory studies could investigate incorporation of colonized microfibers and other microplastics into aggregates, analogous to this study.

Reduced aggregate settling velocity and positive buoyancy of particles alone has the potential to influence marine carbon cycling. Here, we report lower size-specific carbon content of aggregates which, in combination with reduced settling velocity, has the potential to reduce carbon export flux by up to 60% compared to aggregates without microfibers (Fig. 4). This reduction was in the range of 15 to 50% for microfiber concentrations typical of those found in the English Channel and Atlantic Ocean, respectively. We hypothesize that the reduction in carbon flux is likely even higher in situ, as reduced settling velocities would prolong the residence times of microfiber-containing aggregates in the ocean surface and thus extend the time available for microbial remineralization in the upper water column (Ploug et al. 1999). Longer residence times would also increase the probability of encounters with aggregate-feeding zooplankton such as copepods (Poulsen and Kiørboe 2005; Iversen and Poulsen 2007; Lombard et al. 2013), salps (Iversen et al. 2016), polychaetes (Christiansen et al. 2018), and protozoans (Poulsen and Iversen 2008; Poulsen et al. 2011). Additionally, ingestion of microfibers by zooplankton can alter sinking rates of zooplankton faecal pellets (Cole et al. 2016), potentially further exacerbating the effects observed in this study. There is also evidence that the presence of microplastics increases release of chromophoric dissolved organic matter (Galgani et al. 2018), and can affect the water column oxygen inventory due to reduced grazing pressure on primary producers and increased particle remineralization (Kvale et al. 2021), all of which are likely to influence marine carbon dynamics. Together, this would result in higher microbial degradation and turnover of microfiber-containing aggregates in the upper water column where microbially respired CO_2_ will be readily exchanged with the atmosphere. Hence, incorporation of buoyant microfibers into aggregates may substantially reduce the efficiency of the biological carbon pump in a high plastic world. In regional areas which play a pivotal role in the biological carbon pump, e.g., the Southern Ocean (Khatiwala et al. 2013), this could have disproportionate impacts.

**Fig. 4.**
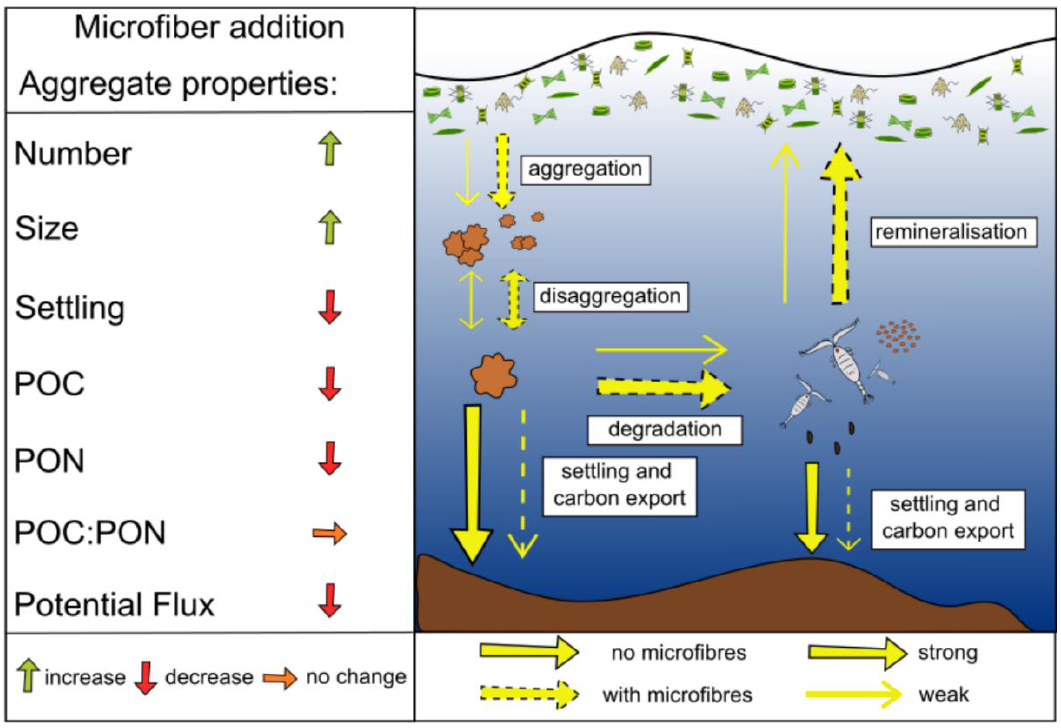
Direction and magnitude of changes in aggregate properties and carbon export related to plastic microfiber incorporation.

Current studies show that all major ocean basins are already heavily contaminated by microplastic (Gago et al. 2018; Peeken et al. 2018; Bergmann et al. 2019). Plastic debris can be found throughout the marine environment (Barnes et al. 2009) and it profoundly affects carbon and nutrient cycling, as well as microbial processes (Galgani and Loiselle 2021, and references therein), including reduced export of zooplankton faecal pellets (Wieczorek et al. 2019). The results of our study, which found reduced size, sinking velocity and carbon content of particles with incorporated microfibers, suggests that the present-day functioning of the biological carbon pump could already be affected by microplastics. Consequently, this may result in ocean services not working at peak efficiency, which has implications for mitigating the increasing atmospheric CO_2_ levels via carbon sequestrations in the deep ocean and sediments. The impact of microplastics and associated debris should be explored in combination with the cascade of impacts arising from increasing atmospheric CO_2,_ and the role this may play in altering the efficiency of the ocean as a carbon sink and wider ecosystem processes.

## Acknowledgements

We thank Christiane Lorentzen for POC and PON analyses. We thank Hannah Marchant for discussions during the writing of the manuscript. CR and CM are funded by the University of Plymouth, CR, CMF, and MHI are funded by the Alfred Wegener Institute Helmholtz Center for Polar and Marine Research and the DFG-Research Center/Cluster of Excellence “The Ocean in the Earth System”: EXC-2077-390741603, AK and JM are funded by the Max Planck Institute for Marine Microbiology. This study was funded by the HGF Young Investigator Group SeaPump “Seasonal and regional food web interactions with the biological pump”: VH-NG-1000.

## Author contribution statement

CR: Conceptualization (equal); Investigation (lead); Writing – original draft (lead); Writing – review and editing (equal); formal analysis (equal); Visualization (equal). CFL: Investigation (supporting); Writing – original draft (lead); writing – review and editing (equal); formal analysis (lead); Visualization (equal). AK: Investigation (lead); Writing – review and editing (equal). JM: Supervision (supporting); writing – review and editing (equal). CBM: Funding Acquisition (lead); writing – review and editing (equal); Supervision (equal). MHI: Conceptualization (equal); Resources (lead); Writing – original draft (supporting); Writing – review and editing (equal); Investigation (supporting); formal analysis (supporting); Project Administration (lead); Supervision (equal).

## Conflict of interest

The authors declare no conflict of interest.

## Data availability statement

The data collected for this study will be made publicly available in the PANGAEA Open Access library.

## Notes

### Competing Interest Statement

The authors have declared no competing interest.

### Summary of Updates

Title revised; author affiliations updated; Figures 1, 2, and S2 revised; discussion revised

## References

Alldredge, A. L., and C. Gotschalk. 1988. In situ settling behavior of marine snow. Limnol. Oceanogr. 33: 339–351. doi:10.4319/lo.1988.33.3.0339

Alldredge, A. L., and M. W. Silver. 1988. Characteristics, dynamics and significance of marine snow. Prog. Oceanogr. 20: 41–82. doi:10.1016/0079-6611(88)90053-5

Amaral-Zettler, L. A., E. R. Zettler, T. J. Mincer, M. A. Klaassen, and S. M. Gallager. 2021. Biofouling impacts on polyethylene density and sinking in coastal waters: A macro/micro tipping point? Water Res. 201: 117289. doi:10.1016/j.watres.2021.117289

Andrady, A. L., and M. A. Neal. 2009. Applications and societal benefits of plastics. Philos. Trans. R. Soc. B Biol. Sci. 364: 1977–1984. doi:10.1098/rstb.2008.0304

Barnes, D. K. A., F. Galgani, R. C. Thompson, and M. Barlaz. 2009. Accumulation and fragmentation of plastic debris in global environments. Philos. Trans. R. Soc. B Biol. Sci. 364: 1985–1998. doi:10.1098/rstb.2008.0205

Bergmann, M., S. Mützel, S. Primpke, M. B. Tekman, J. Trachsel, and G. Gerdts. 2019. White and wonderful? Microplastics prevail in snow from the Alps to the Arctic. Sci. Adv. 5. doi:10.1126/sciadv.aax1157

Boyd, P. W., H. Claustre, M. Levy, D. A. Siegel, and T. Weber. 2019. Multi-faceted particle pumps drive carbon sequestration in the ocean. Nature 568: 327–335. doi:10.1038/s41586-019-1098-2

Browne, M. A., P. Crump, S. J. Niven, E. Teuten, A. Tonkin, T. Galloway, and R. Thompson. 2011. Accumulation of Microplastic on Shorelines Woldwide: Sources and Sinks. Environ. Sci. Technol. 45: 9175–9179. doi:10.1021/es201811s

Carbery, M., W. O’Connor, and T. Palanisami. 2018. Trophic transfer of microplastics and mixed contaminants in the marine food web and implications for human health. Environ. Int. 115: 400–409. doi:10.1016/j.envint.2018.03.007

Christiansen, S., H.-J. Hoving, F. Schütte, and others. 2018. Particulate matter flux interception in oceanic mesoscale eddies by the polychaete Poeobius sp. Limnol. Oceanogr. 63: 2093–2109. doi:10.1002/lno.10926

Cole, M. C., P. Lindeque, E. S. Fileman, J. Clark, C. Lewis, C. Halsband, and T. S. Galloway. 2016. Microplastics Alter the Properties and Sinking Rates of Zooplankton Faecal Pellets. Environ. Sci. Technol. 50: 3239–3246. doi:10.1021/acs.est.5b05905

Cole, M., H. Webb, P. K. Lindeque, E. S. Fileman, C. Halsband, and T. S. Galloway. 2014. Isolation of microplastics in biota-rich seawater samples and marine organisms. Sci. Rep. 4: 4528. doi:10.1038/srep04528

Gago, J., O. Carretero, A. V. Filgueiras, and L. Viñas. 2018. Synthetic microfibers in the marine environment: A review on their occurrence in seawater and sediments. Mar. Pollut. Bull. 127: 365–376. doi:10.1016/j.marpolbul.2017.11.070

Galgani, L., A. Engel, C. Rossi, A. Donati, and S. A. Loiselle. 2018. Polystyrene microplastics increase microbial release of marine Chromophoric Dissolved Organic Matter in microcosm experiments. Sci. Rep. 8: 14635. doi:10.1038/s41598-018-32805-4

Galgani, L., I. Goßmann, B. Scholz-Böttcher, X. Jiang, Z. Liu, L. Scheidemann, C. Schlundt, and A. Engel. 2022. Hitchhiking into the Deep: How Microplastic Particles are Exported through the Biological Carbon Pump in the North Atlantic Ocean. Environ. Sci. Technol. 56: 15638–15649. doi:10.1021/acs.est.2c04712

Galgani, L., and S. A. Loiselle. 2021. Plastic pollution impacts on marine carbon biogeochemistry. Environ. Pollut. Barking Essex 1987 268: 115598. doi:10.1016/j.envpol.2020.115598

Geyer, R., J. R. Jambeck, and K. L. Law. 2017. Production, use, and fate of all plastics ever made. Sci. Adv. 3: e1700782. doi:10.1126/sciadv.1700782

Gregory, M. R. 1996. Plastic ‘scrubbers’ in hand cleansers: a further (and minor) source for marine pollution identified. Mar. Pollut. Bull. 32: 867–871. doi:10.1016/S0025-326X(96)00047-1

Guillard, R. R. L. 1975. Culture of Phytoplankton for Feeding Marine Invertebrates, p. 29–60. In W.L. Smith and M.H. Chanley [eds.], Culture of Marine Invertebrate Animals: Proceedings — 1st Conference on Culture of Marine Invertebrate Animals Greenport. Springer US.

Iversen, M. H., and H. Ploug. 2010. Ballast minerals and the sinking carbon flux in the ocean: carbon-specific respiration rates and sinking velocity of marine snow aggregates. Biogeosciences 7: 2613–2624. doi:10.5194/bg-7-2613-2010

Iversen, M. H., and L. K. Poulsen. 2007. Coprorhexy, coprophagy, and coprochaly in the copepods Calanus helgolandicus, Pseudocalanus elongatus, and Oithona similis. Mar. Ecol. Prog. Ser. 350: 79–89. doi:10.3354/meps07095

Iversen, M., E. Pakhomov, B. Hunt, H. van der Jagt, D. Wolf-Gladrow, and C. Klaas. 2016. Sinkers or floaters? Contribution from salp pellets to the export flux during a large bloom event in the Southern Ocean. Deep Sea Res. Part II Top. Stud. Oceanogr. 138. doi:10.1016/j.dsr2.2016.12.004

Jambeck, J. R., R. Geyer, C. Wilcox, T. R. Siegler, M. Perryman, A. Andrady, R. Narayan, and K. L. Law. 2015. Plastic waste inputs from land into the ocean. Science 347: 768–771. doi:10.1126/science.1260352

Kaiser, D., N. Kowalski, and J. J. Waniek. 2017. Effects of biofouling on the sinking behavior of microplastics. Environ. Res. Lett. 12: 124003. doi:10.1088/1748-9326/aa8e8b

Kanhai, L. D. K., R. Officer, O. Lyashevska, R. C. Thompson, and I. O’Connor. 2017. Microplastic abundance, distribution and composition along a latitudinal gradient in the Atlantic Ocean. Mar. Pollut. Bull. 115: 307–314. doi:10.1016/j.marpolbul.2016.12.025

Khatiwala, S., T. Tanhua, S. Mikaloff Fletcher, and others. 2013. Global ocean storage of anthropogenic carbon. Biogeosciences 10: 2169–2191. doi:10.5194/bg-10-2169-2013

Kvale, K., A. E. F. Prowe, C.-T. Chien, A. Landolfi, and A. Oschlies. 2020. The global biological microplastic particle sink. Sci. Rep. 10: 16670. doi:10.1038/s41598-020-72898-4

Kvale, K., A. E. F. Prowe, C.-T. Chien, A. Landolfi, and A. Oschlies. 2021. Zooplankton grazing of microplastic can accelerate global loss of ocean oxygen. Nat. Commun. 12: 2358. doi:10.1038/s41467-021-22554-w

Lepot, L., K. De Wael, F. Gason, and B. Gilbert. 2008. Application of Raman spectroscopy to forensic fibre cases. Sci. Justice 48: 109–117. doi:10.1016/j.scijus.2007.09.013

Liu, J., Q. Liu, L. An, and others. 2022. Microfiber Pollution in the Earth System. Rev. Environ. Contam. Toxicol. 260: 13. doi:10.1007/s44169-022-00015-9

Lombard, F., M. Koski, and T. Kiørboe. 2013. Copepods use chemical trails to find sinking marine snow aggregates. Limnol. Oceanogr. 58: 185–192. doi:10.4319/lo.2013.58.1.0185

Long, M., B. Moriceau, M. Gallinari, C. Lambert, A. Huvet, J. Raffray, and P. Soudant. 2015. Interactions between microplastics and phytoplankton aggregates: Impact on their respective fates. Mar. Chem. 175: 39–46. doi:10.1016/j.marchem.2015.04.003

Michels, J., A. Stippkugel, M. Lenz, K. Wirtz, and A. Engel. 2018. Rapid aggregation of biofilm-covered microplastics with marine biogenic particles. Proc. R. Soc. B Biol. Sci. 285: 20181203. doi:10.1098/rspb.2018.1203

Peeken, I., S. Primpke, B. Beyer, and others. 2018. Arctic sea ice is an important temporal sink and means of transport for microplastic. Nat. Commun. 9: 1505. doi:10.1038/s41467-018-03825-5

Pirc, U., M. Vidmar, A. Mozer, and A. Kržan. 2016. Emissions of microplastic fibers from microfiber fleece during domestic washing. Environ. Sci. Pollut. Res. 23: 22206–22211. doi:10.1007/s11356-016-7703-0

Ploug, H., H. P. Grossart, F. Azam, and B. B. Jørgensen. 1999. Photosynthesis, respiration, and carbon turnover in sinking marine snow from surface waters of Southern California Bight: Implications for the carbon cycle in the ocean. Mar. Ecol.-Prog. Ser. 179: 1–11.

Ploug, H., M. H. Iversen, and G. Fischer. 2008. Ballast, sinking velocity, and apparent diffusivity within marine snow and zooplankton fecal pellets: Implications for substrate turnover by attached bacteria. Limnol. Oceanogr. 53: 1878–1886. doi:10.4319/lo.2008.53.5.1878

Ploug, H., A. Terbrüggen, A. Kaufmann, D. Wolf-Gladrow, and U. Passow. 2010. A novel method to measure particle sinking velocity in vitro, and its comparison to three other in vitro methods. Limnol. Oceanogr. Methods 8: 386–393.

Porter, A., B. P. Lyons, T. S. Galloway, and C. Lewis. 2018. Role of Marine Snows in Microplastic Fate and Bioavailability. Environ. Sci. Technol. 52: 7111–7119. doi:10.1021/acs.est.8b01000

Poulsen, L. K., and M. H. Iversen. 2008. Degradation of copepod fecal pellets: key role of protozooplankton. Mar. Ecol. Prog. Ser. 367: 1–13. doi:10.3354/meps07611

Poulsen, L. K., and T. Kiørboe. 2005. Coprophagy and coprorhexy in the copepods Acartia tonsa and Temora longicornis: clearance rates and feeding behaviour. Mar. Ecol. Prog. Ser. 299: 217–227. doi:10.3354/meps299217

Poulsen, L. K., M. Moldrup, T. Berge, and P. J. Hansen. 2011. Feeding on copepod fecal pellets: a new trophic role of dinoflagellates as detritivores. Mar. Ecol. Prog. Ser. 441: 65–78. doi:10.3354/meps09357

Ryan, P. G., and C. L. Moloney. 1993. Marine litter keeps increasing. Nature 361: 23–23. doi:10.1038/361023a0

Ryan, P. G., C. J. Moore, J. A. van Franeker, and C. L. Moloney. 2009. Monitoring the abundance of plastic debris in the marine environment. Philos. Trans. R. Soc. B Biol. Sci. 364: 1999–2012. doi:10.1098/rstb.2008.0207

Setälä, O., M. Lehtiniemi, R. Coppock, and M. Cole. 2018. Chapter 11 - Microplastics in Marine Food Webs, p. 339–363. In E.Y. Zeng [ed.], Microplastic Contamination in Aquatic Environments. Elsevier.

Shore, E. A., J. A. deMayo, and M. H. Pespeni. 2021. Microplastics reduce net population growth and fecal pellet sinking rates for the marine copepod, Acartia tonsa. Environ. Pollut. 284: 117379. doi:10.1016/j.envpol.2021.117379

Taylor, M. L., C. Gwinnett, L. F. Robinson, and L. C. Woodall. 2016. Plastic microfibre ingestion by deep-sea organisms. Sci. Rep. 6: 33997. doi:10.1038/srep33997

Thompson, R. C., Y. Olsen, R. P. Mitchell, A. Davis, S. J. Rowland, A. W. G. John, D. McGonigle, and A. E. Russell. 2004. Lost at Sea:Where Is All the Plastic? Science 304: 838–838. doi:10.1126/science.1094559

Vaksmaa, A., M. Egger, C. Lüke, P. D. Martins, R. Rosselli, A. A. Asbun, and H. Niemann. 2022. Microbial communities on plastic particles in surface waters differ from subsurface waters of the North Pacific Subtropical Gyre. Mar. Pollut. Bull. 182: 113949. doi:10.1016/j.marpolbul.2022.113949

Van Cauwenberghe, L., A. Vanreusel, J. Mees, and C. R. Janssen. 2013. Microplastic pollution in deep-sea sediments. Environ. Pollut. 182: 495–499. doi:10.1016/j.envpol.2013.08.013

Volk, T., and M. I. Hoffert. 1985. Ocean Carbon Pumps: Analysis of Relative Strengths and Efficiencies in Ocean-Driven Atmospheric CO2 Changes, p. 99–110. In E.T. Sundquist and W.S. Broecker [eds.], The Carbon Cycle and Atmospheric CO2: Natural Variations Archean to Present. American Geophysical Union.

Wieczorek, A. M., P. L. Croot, F. Lombard, J. N. Sheahan, and T. K. Doyle. 2019. Microplastic Ingestion by Gelatinous Zooplankton May Lower Efficiency of the Biological Pump. Environ. Sci. Technol. 53: 5387–5395. doi:10.1021/acs.est.8b07174

Woodall, L. C., A. Sanchez-Vidal, M. Canals, and others. 2014. The deep sea is a major sink for microplastic debris. R. Soc. Open Sci. 1: 140317. doi:10.1098/rsos.140317

Zettler, E. R., T. J. Mincer, and L. A. Amaral-Zettler. 2013. Life in the “plastisphere”: microbial communities on plastic marine debris. Environ. Sci. Technol. 47: 7137–7146. doi:10.1021/es401288x

Zhao, S., J. E. Ward, M. Danley, and T. J. Mincer. 2018. Field-Based Evidence for Microplastic in Marine Aggregates and Mussels: Implications for Trophic Transfer. Environ. Sci. Technol. 52: 11038–11048. doi:10.1021/acs.est.8b03467

