## Supporting information for "Microplastics may reduce the efficiency of the biological carbon pump by decreasing the settling velocity and carbon content of marine snow"

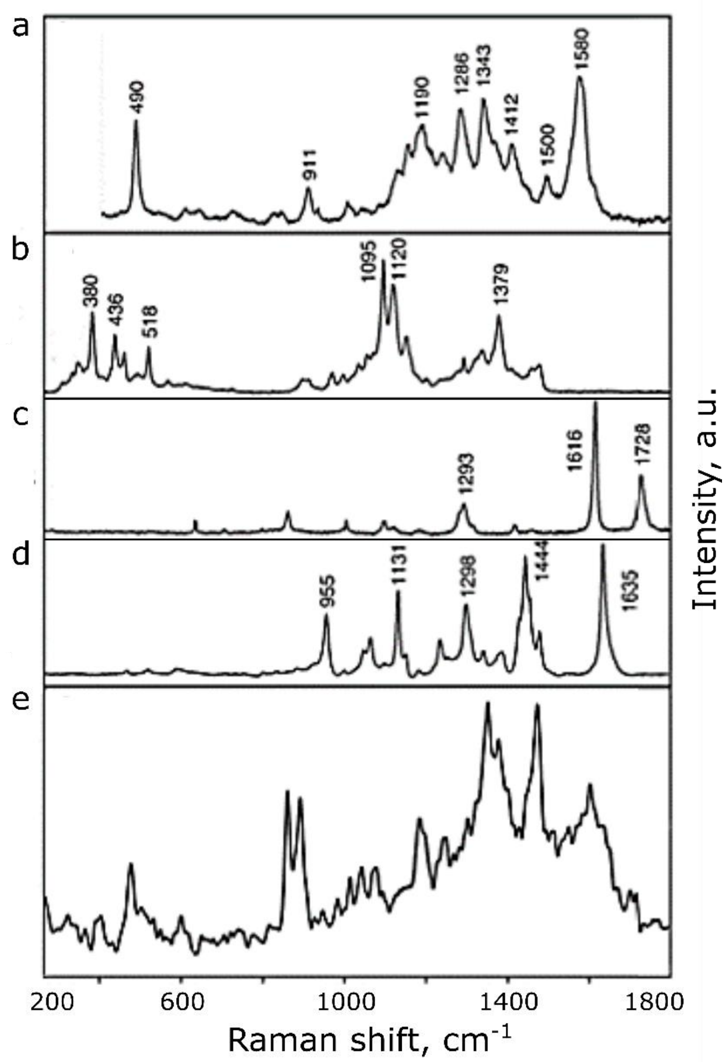


**Fig. S1.** Raman spectra of microfibers used in experiments: dyed cotton fiber (a), undyed cotton (b), PET (c), polyamide (d) and microfiber sample (e).

**Table S1.** Microfiber concentration at $t_0$, aggregate number and aggregate microfiber, POC, and PON content at $t_{fin}$, predicted POC flux to 100 m depth, and decrease in POC flux for each treatment.

| **Treatment** | **Control** | **Low** | **Medium** | **High** |
|---|---|---|---|---|
| Microfiber concentration at $t_0$ ($L^{-1}$) | 0 | 240 (±0.2) | 680 (±0.9) | 840 (±1.1) |
| Aggregates at $t_{fin}$ (n) | 7 | 14 | 18 | 13 |
| Aggregate microfiber conc. at $t_{fin}$ ($mm^{-3}$) | 0 | 0.026 (±0.0004) | 0.057 (±0.01) | 0.133 (±0.02) |
| POC (µgC $mm^{-3}$) | 2.43 | 1.88 | 2.00 | 1.26 |
| PON (µgN $mm^{-3}$) | 0.32 | 0.26 | 0.24 | 0.16 |
| Predicted flux to 100 m (gC $m^{-2}$ $d^{-1}$) | 60.2 (±21.1) | 51.7 (±9.1) | 39.6 (±3.5) | 29.7 (±2.8) |
| % Decrease in flux compared to control | - | 14 | 34 | 51 |

**Table S2.** ANOVA results for influence of different microfiber treatments on aggregate volume, aggregate POC, aggregate flux and number of plastics incorporated on 3-day average POC flux.

| **Parameter** | **Degrees of freedom** | **P-value** | **Tukey's HSD significant pairwise comparisons** |
|---|---|---|---|
| Aggregate Volume | 3 | ns | - |
| Aggregate POC | 3 | 0.02* | High – Control<br>High – Low |
| $POC_{100}$ flux | 3 | ns | - |
| Microfiber content vs $POC_{100}$ flux | 3 | <0.01 *** | Medium – Control<br>High – Control<br>Medium – Low<br>High – Low<br>High – Medium |

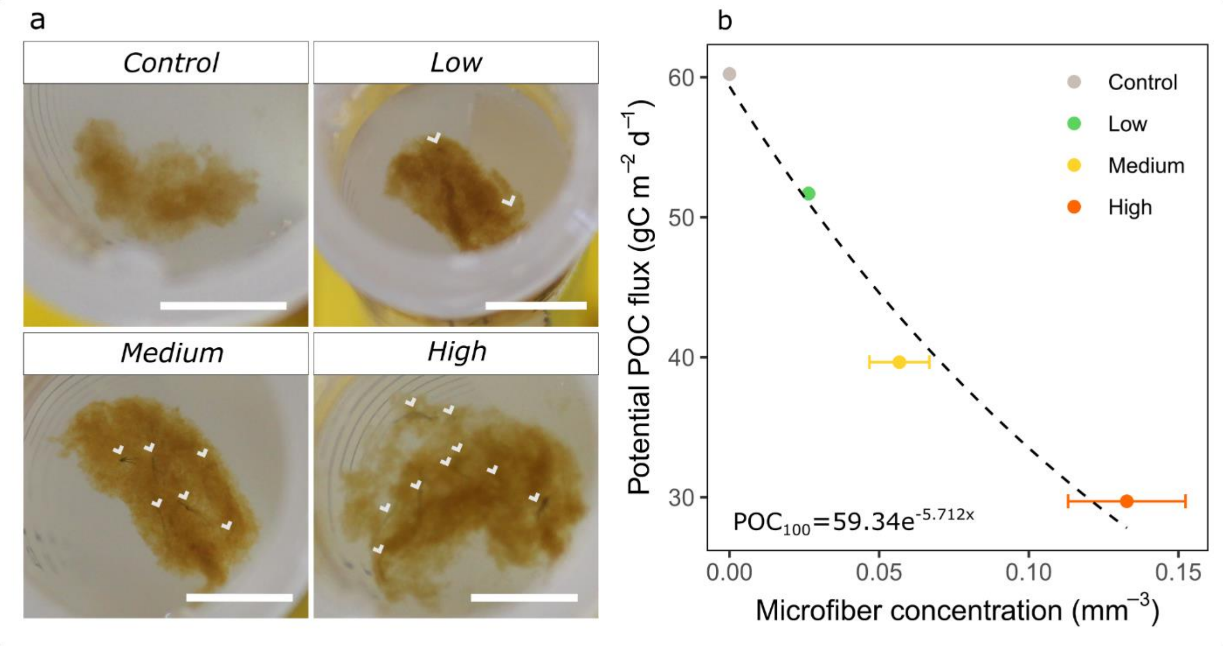


**Fig. S2.** Examples of aggregates from each treatment with microfibers incorporated into the matrix (white arrows); scale bar=0.5 cm (a). Correlation between potential POC flux to 100 m and number of microfibers per aggregate volume, averaged over days 5-7 for each treatment (b). Microfiber concentration is highly statistically significant between treatments (Table S2).
